# Intricate regulation of ribosome biogenesis genes in response to mTORC1 signaling

**DOI:** 10.1101/2021.01.04.425249

**Authors:** Sanjay Kumar, Muneera Mashkoor, Priya Balamurugan, Anne Grove

## Abstract

Genes encoding ribosomal proteins are repressed in response to inhibition of mTORC1. In *Saccharomyces cerevisiae*, this involves dissociation of the activator Ifh1p in a process that depends on Utp22p, a protein that also functions in pre-rRNA processing. Ifh1p has a paralog, Crf1p, which can mediate mTORC1 inhibition by acting as a repressor. Ifh1p and Crf1p derive from a common ancestor, which may have acted as both an activator and a repressor. We report here that *UTP22* and *RRP7*, which encodes another pre-rRNA processing factor, are controlled by mTORC1; both gene promoters are bound by Ifh1p, which dissociates on mTORC1 inhibition. Notably, Crf1p acts as an activator as evidenced by reduced expression in a *crf1Δ* strain. By contrast, Crf1p is required to repress expression of *HMO1*, which encodes a cofactor involved in communicating mTORC1 activity to target genes. Our data therefore indicate that Crf1p exhibits the dual repressor/activator functions of the Ifh1p-Crf1p ancestor.

## Introduction

The evolutionarily conserved mechanistic Target of Rapamycin Complex 1 (mTORC1) senses nutrient sufficiency and cellular stress to control cell growth and ribosome biogenesis, the latter requiring significant cellular energy. In budding yeast (*Saccharomyces cerevisiae*), the multisubunit mTORC1 contains either of the two homologous phosphatidylinositol 3-kinase-related subunits, Tor1p or Tor2p, and its inhibition under nutrient depletion or stress is relayed through several direct or indirect targets (Gonzalez and Hall, 2017; Laribee, 2018; Loewith et al., 2002; Sabatini, 2017; Saxton and Sabatini, 2017). Amino acid sufficiency is detected by cytoplasmic mTORC1 localized to the vacuolar surface, resulting in phosphorylation of mTORC1 targets (Binda et al., 2009; Sancak et al., 2008). The best characterized mTORC1 direct targets include regulators of translation such as Sch9p (the yeast ortholog of mammalian S6K1) and 4E-BP1 (Hara et al., 1998; Loewith and Hall, 2011; Urban et al., 2007). Treatment with the macrolide rapamycin phenocopies nutrient deprivation (Heitman et al., 1991).

The mTORC1 complex also exerts its functions in the nucleus where it regulates ribosome biogenesis. Specifically, the coordination of rRNA production and ribosomal protein (RP) synthesis is intricately coupled to nutrient and stress signaling pathways (Xiao and Grove, 2009). In yeast, ∼2,000 ribosomes are assembled per minute during normal growth, and a fine balance is required to regulate expression of the RNA Polymerase (Pol) I-transcribed polycistronic rRNA, the Pol III-transcribed 5S rRNA, and the 138 Pol II-transcribed RP genes as well as genes encoding proteins involved in ribosome biogenesis (RiBi) (Woolford and Baserga, 2013). That coordination of transcriptional output in response to metabolic resources is critical is evidenced by the proteotoxic stress imposed by dysregulation (Hein et al., 2013; James et al., 2014; Tye et al., 2019).

A direct role for mTORC1 in transcriptional control of rRNA genes is indicated by recruitment of both yeast Tor1p and mammalian Tor kinases to the rDNA in a nutrient-dependent fashion (Li et al., 2006; Tsang et al., 2010). Transcription of the rDNA also depends on high mobility group (HMG) proteins (Vizoso-Vazquez et al., 2018); in mammals, the Upstream Binding Factor (UBF) is a target for S6K1 (Hannan et al., 2003), and in yeast, Hmo1p has been shown to bind the rDNA and to promote transcription in an mTORC1-dependent manner (Gadal et al., 2002; Merz et al., 2008). For yeast 5S rRNA and tRNA genes, transcription depends on mTORC1-mediated phosphorylation of Sch9p, which in turn phosphorylates the repressor Maf1p and keeps it in the cytoplasm (Lee et al., 2009; Vannini et al., 2010; Wei et al., 2009); in mammals, Tor is recruited to the Pol III-transcribed genes, where it directly phosphorylates Maf1 (Michels et al., 2010). Yeast Hmo1p has also been reported to bind the 5S rDNA, albeit at a lower level compared to the 35S rDNA (Hall et al., 2006).

On yeast RP genes, a primary mechanism by which mTORC1 controls expression depends on the constitutively bound Fhl1p (forkhead-like protein), whose binding to gene promoters may be enhanced by simultaneous binding of Hmo1p (Hall et al., 2006). Fhl1p in turn recruits the coactivator Ifh1p (interacts with forkhead-like protein). Inhibition of mTORC1 leads to Yak1p-mediated phosphorylation of the cytoplasmically localized corepressor Crf1p, which then translocates to the nucleus and competes with Ifh1p for binding to Fhl1p to repress RP gene transcription (Jorgensen et al., 2004; Martin et al., 2004; Schawalder et al., 2004; Wade et al., 2004; Zhao et al., 2006). Ifh1p and Crf1p have been reported to be paralogs that originate from an ancestral gene following a whole genome duplication event (Wapinski et al., 2010), and their phosphorylation by Casein Kinase 2 (CK2) was reported to be important for interaction with Fhl1p (Kim and Hahn, 2016). While most RP gene promoters bind both Fhl1p, Ifh1p, and the transcription factor Rap1p, binding of Hmo1p is variable (Kasahara et al., 2007; Knight et al., 2014; Reja et al., 2015). Notably, the participation of Crf1p may also be dependent on strain background as it has been reported not to be required for repression of RP gene activity in the yeast strain W303 (Zhao et al., 2006). In addition, the split-finger protein Sfp1p has been implicated in control of RP genes in response to stress and nutrient limitation (Jorgensen et al., 2004; Marion et al., 2004), and Sfp1p has been detected on RP gene promoters and at promoters driving expression of RiBi genes (Albert et al., 2019; Reja et al., 2015). Another mTORC1 target is the *HMO1* gene, for which mRNA levels decline upon mTORC1 inhibition by rapamycin treatment or by induction of DNA damage stress (Panday et al., 2017; Panday et al., 2015; Xiao et al., 2011). Ifh1p also binds the *HMO1* promoter and, similar to RP genes, it dissociates rapidly upon addition of rapamycin and is replaced with Crf1p (Panday et al., 2017).

The stable release of Ifh1p in response to mTORC1 inhibition requires Utp22p and it is followed by assembly of a multi-subunit protein complex named CURI (Albert et al., 2016). CURI complex formation requires the movement of Ifh1p into the nucleolus and its binding to a pre-existing complex, UTP-C 90S, which is involved in pre-ribosomal RNA processing (Albert et al., 2016). The UTP-C 90S pre-ribosome subcomplex includes CK2, Utp22p and Rrp7p, proteins that are conserved from yeast to mammals (Kornprobst et al., 2016). Thus, “molecular kidnapping” of Ifh1p generates the CURI complex, to which Fhl1p may also be loosely associated, a process that simultaneously sequesters both Ifh1p and components required for pre-ribosomal processing, thereby suppressing ribosome production (Albert et al., 2016; Rudra et al., 2007; Rudra and Warner, 2016).

We show here that mRNA levels of *UTP22* and *RRP7* declined on inhibition of mTORC1. These reduced mRNA levels correlated with dissociation of Ifh1p followed by recruitment of Crf1p to the promoter regions, suggesting that activity of mTORC1 is communicated through Fhl1p and its cofactors. Tor1p and Crf1p were required to communicate mTORC1 inhibition on the *HMO1* gene, but not on *UTP22* and *RRP7*. Notably, optimal *UTP22, RRP7*, and *HMO1* expression during balanced growth required Crf1p. Our data therefore indicate that Crf1p functions as an activator for these genes, while it is required for repression of *HMO1* in response to mTORC1 inhibition.

## Results

### Inhibition of mTORC1 leads to reduced *HMO1, UTP22*, and *RRP7* expression

Inhibition of mTORC1 leads to reduced expression of RP genes as well as *HMO1* by a mechanism that involves Fhl1p and its associated cofactors, Ifh1p and Crf1p. The observation that Ifh1p also associates with the *UTP22* gene raises the possibility that *UTP22* expression is likewise controlled by mTORC1. Since Utp22p and Rrp7p function in pre-rRNA processing, we therefore investigated expression of both *UTP22* and *RRP7*. Upon inhibition of mTORC1 by rapamycin in wildtype DDY3 (which is isogenic to W303), we observed a time-dependent reduction in *UTP22* mRNA abundance, with ∼90% reduction after 1 h of rapamycin treatment (Figure 1A). For *RRP7*, a similar time-dependent reduction in mRNA levels was observed with ∼70% reduction 1 h after addition of rapamycin. This observation suggests an even more sensitive regulation of *UTP22* and *RRP7* on inhibition of mTORC1 by comparison to *HMO1*, for which mRNA abundance levels were reduced ∼30% response to 1 h of rapamycin treatment (Figure 1A).

**Figure 1.**
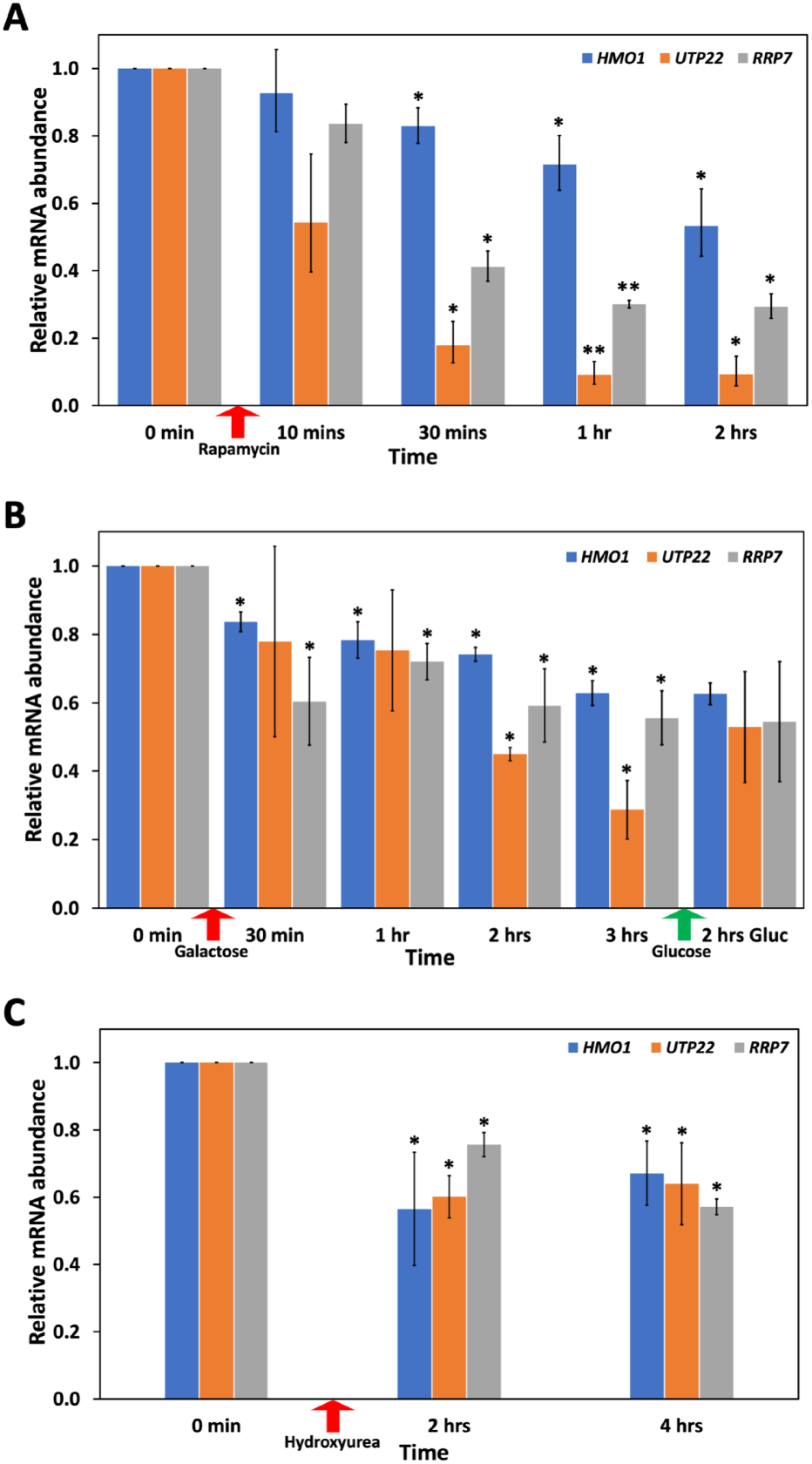
Decreased expression of *UTP22* and *RRP7* in response to DNA damage stress and inhibition of mTORC1. A. Relative abundance of *HMO1* (blue), *UTP22* (orange), and *RRP7* (grey) mRNA before and at the indicated times after administration of rapamycin (red arrow). B. mRNA abundance before and after administration of galactose to induce DSB formation at the *MAT* locus (red arrow) or addition of glucose to terminate expression of HO endonuclease (green arrow). C. mRNA abundance before and after exposure of cultures to hydroxyurea (red arrow). Error bars represent standard deviation from three biological replicates. Transcript levels were calculated using 2^-ΔΔC^_T_ relative to the reference gene and reported relative to the corresponding unsupplemented cultures (0 min). Asterisks denote statistically significant differences compared to unsupplemented cultures based on a Student’s t-test; *, p<0.05; **, p<0.001.

### DNA double strand break and replication stress also reduces expression of *HMO1, UTP22*, and *RRP7*

Using a plasmid-encoded HO endonuclease under control of the *GAL* promoter to site-specifically introduce a DNA double strand break (DSB) at the mating type (*MAT*) locus, we previously reported reduced *HMO1* mRNA abundance upon DSB induction (Panday et al., 2017). We therefore tested the effect of DSB induction on *UTP22* and *RRP7* mRNA levels. A reduction in *UTP22* mRNA abundance was observed, particularly after more than 2 h of DSB induction, with mRNA levels reaching ∼30% by 3 h as compared to the initial level without the DSB (Figure 1B). Stopping the break induction by adding glucose and allowing repair, resulted in a trend towards increased *UTP22* mRNA levels. A similar trend of time-dependent reduction in *RRP7* mRNA level was detectable 30 min post DSB induction and reaching ∼55% after 3 h when compared to cells without DSB. There was no increase in *RRP7* mRNA levels by 2 h after addition of glucose to terminate HO endonuclease expression. These observations were comparable to what has been reported earlier in case of *HMO1* (Figure 1B).

We also tested the response of our genes of interest upon replication stress induced by hydroxyurea treatment for up to 4 hours. We observed a reduction in *HMO1* mRNA level of ∼30% four hours post HU addition, with a similar decrease in *UTP22* and *RRP7* mRNA levels after HU treatment (Figure 1C).

### Repression of *HMO1, UTP22*, and *RRP7* correlates with dissociation of Ifh1p

As noted above, transcription factors Hmo1p, Fhl1p, Ifh1p, and Crf1p have been implicated in mTORC1-mediated control of RP and *HMO1* genes. Using Chromatin Immunoprecipitation (ChIP) followed by qPCR and yeast strains expressing 3X-FLAG-tagged Ifh1p, Crf1p, Hmo1p, or Fhl1p, we tested binding of these proteins to the *UTP22* and *RRP7* genes. As reported previously (Panday et al., 2017), Ifh1p dissociates rapidly (<5 min) from the *HMO1* promoter (Figure S1A) after administration of rapamycin and is barely detectable after 30 min (Figure S1B). The dissociation of Ifh1p is followed by enhanced binding of Crf1p ∼15 min post rapamycin treatment as reported earlier, and Crf1 remained bound after 30 minutes of rapamycin treatment (Figure S1C). We also confirmed the dissociation of Hmo1p from its own promoter after 2 h of rapamycin treatment, whereas Fhl1p binding after 2 h was not significantly reduced (Figure S1D-E). These binding signals were higher for amplicons near the IFHL site; this site represents the binding site for Fhl1p, as confirmed by ChIP and by the observation that mutagenesis attenuates the *HMO1* promoter response to rapamycin (Xiao et al., 2011).

We used FIMO (Find Individual Motif Occurrences), which is part of the MEME suite (http://meme-suite.org/) to predict DNA sequence motifs in *UTP22* and *RRP7* gene promoters by analyzing 1 KB upstream of *HMO1, UTP22*, and *RRP7* along with the same length of RP gene promoters (Grant et al., 2011). We observed the consensus DNA sequence motif representing the IFHL site in *HMO1, UTP22*, and *RRP7* at the positions noted in Figures S1, 2, and 3. On *UTP22*, binding of Ifh1p was detected near the predicted IFHL site, followed by significant dissociation after 30 min of rapamycin treatment (Figure 2A). Enhanced Crf1p binding was observed following rapamycin treatment, however, Crf1p was detectable at the IFHL site even prior to addition of rapamycin (Figure 2C). For *RRP7*, Ifh1p dissociated following rapamycin treatment and was replaced by Crf1p (Figure 3A,B). We also observed a marked dissociation of Hmo1p from *UTP22* and *RRP7* after 2 h of mTORC1 inhibition by rapamycin and no significant loss of Fhl1p binding from the respective sites (Figures 2 and 3). This pattern of transcription factor binding in response to mTORC1 inhibition largely mirrors that observed for the *HMO1* and RP genes, with the possible exception of detectable Crf1p binding to *UTP22* prior to mTORC1 inhibition, and it suggests that mTORC1 inhibition is communicated to the *UTP22* and *RRP7* genes via Fhl1p and its cofactors.

**Figure 2.**
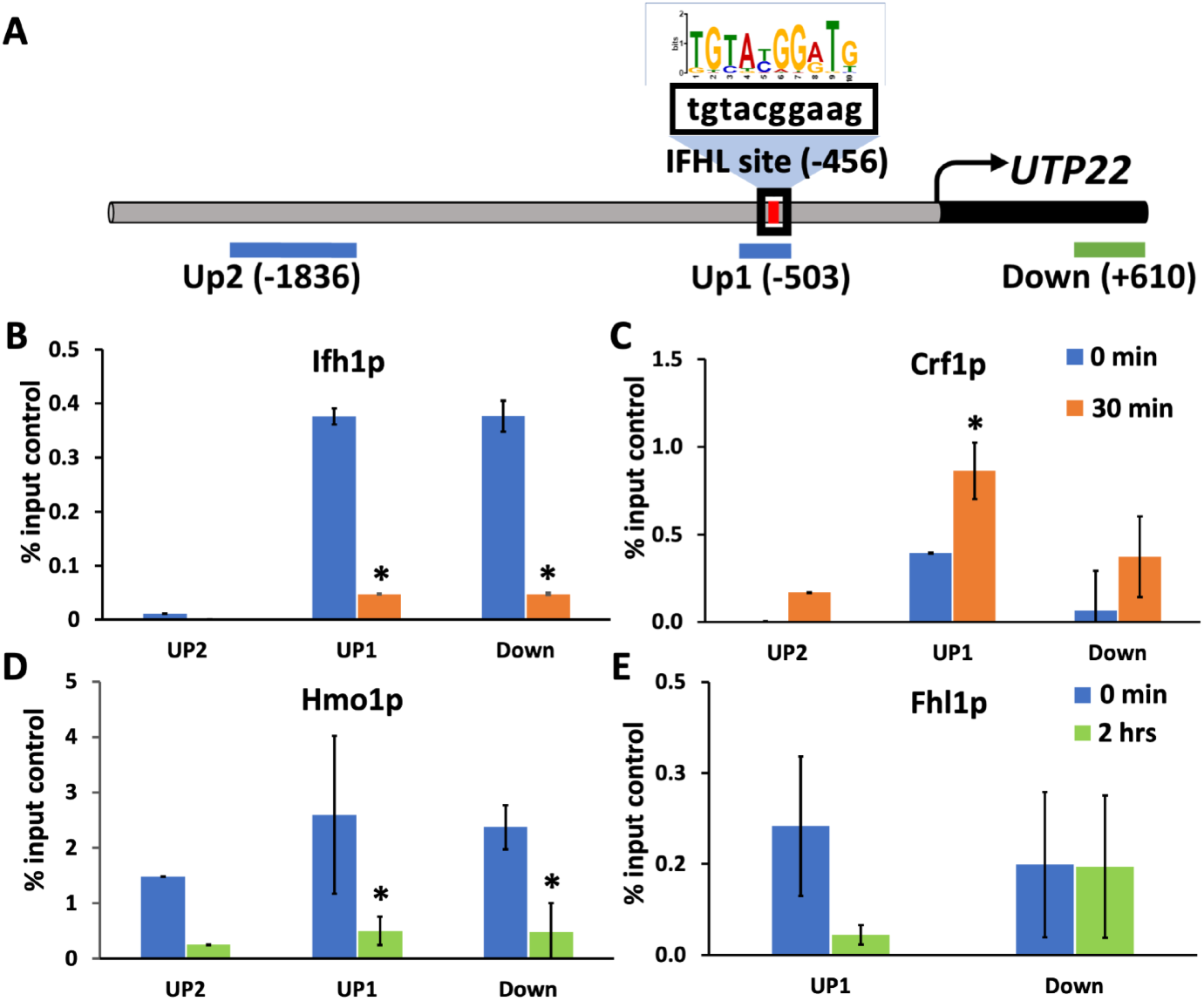
Transcription factor binding to the *UTP22* gene. A. *UTP22* upstream region with predicted IFHL site shown in red and expanded to show sequence; consensus IFHL site shown above. Positions of amplicons are identified, with numbers representing the upstream edge. B-E. Binding of Ifh1p, Crf1p, Hmo1p, and Fhl1p at the indicated positions as determined by ChIP using antibodies to FLAG-tagged proteins; amplicons are identified at the top. Blue bars represent binding before addition of rapamycin, while orange and green bars represent binding detected 30 min and 2 h following addition of rapamycin, respectively. Data are normalized to the corresponding input control and are presented as the average of three biological replicates; error bars represent SD.

**Figure 3.**
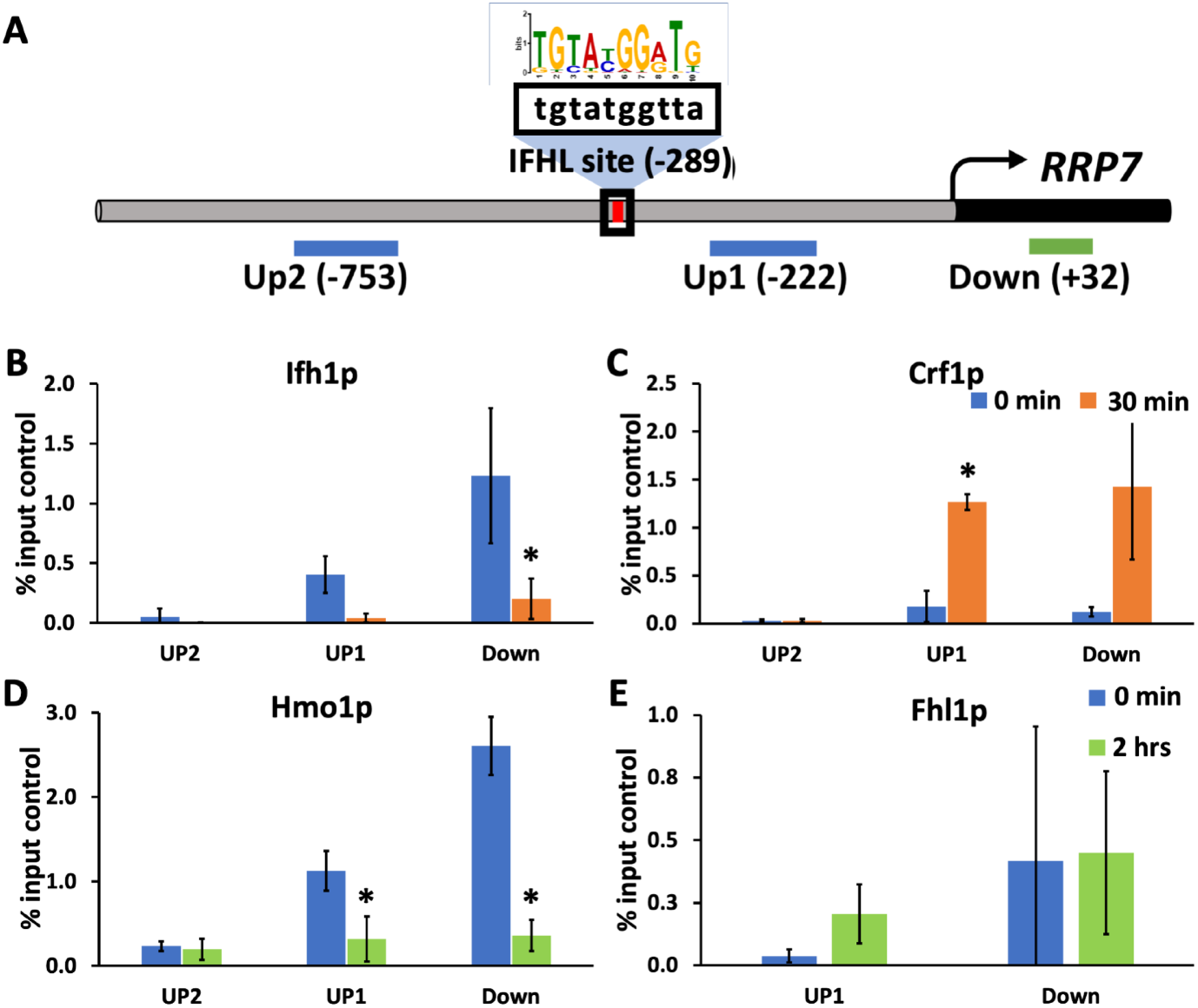
Transcription factor binding to the *RRP7* gene. A. *RRP7* upstream region with IFHL site shown in red and expanded to show sequence; consensus IFHL site shown above. Positions of amplicons are identified, with numbers representing the upstream edge. B-E. Binding of Ifh1p, Crf1p, Hmo1p, and Fhl1p at the indicated positions as determined by ChIP using antibodies to FLAG-tagged proteins; amplicons are identified at the top. Blue bars represent binding before addition of rapamycin, while orange and green bars represent binding detected 30 min and 2 h following addition of rapamycin, respectively. Data are normalized to the corresponding input control and are presented as the average of three biological replicates; error bars represent SD.

### Differential Crf1p requirement for regulation of *HMO1, UTP22*, and *RRP7*

A reduced mRNA abundance was observed for *HMO1, UTP22*, and *RRP7* upon rapamycin treatment (Figure 1A) with an accompanying reduction in Ifh1p binding and recruitment of Crf1p at their promoter regions. To determine if Crf1p actively participates in the rapamycin-induced change in mRNA abundance, we used a *crf1Δ* strain. While Crf1p was reported to mediate repression of RP genes upon mTORC1 inhibition, this effect was not seen in W303 (Zhao et al., 2006), which is the strain used in these experiments. Interestingly, deletion of *CRF1* resulted in *HMO1* mRNA levels to remain high despite inhibition of mTORC1 until 1 h after rapamycin addition. The mRNA levels were slightly elevated 30 min after rapamycin addition and then reduced to ∼60% after 2 h of rapamycin treatment (Figure 4A). In contrast, a consistent reduction in *UTP22* and *RRP7* mRNA levels was observed, reaching 30-40% after 1 h. This indicates that Crf1p is not required for rapamycin-mediated reduction in *UTP2*2 and *RRP7* mRNA accumulation, but that it is essential for regulation of *HMO1* expression. Consistent with the observation that repression of RP gene expression is independent of Crf1p in W303 (DDY3) cells and with the interpretation that cytoplasmic mTORC1 is sufficient for regulation of RP gene expression (Li et al., 2006; Zhao et al., 2006), deletion of *CRF1* or *TOR1* had no effect on rapamycin-mediated reduction in expression of select RP genes (Figure S2A-C).

**Figure 4.**
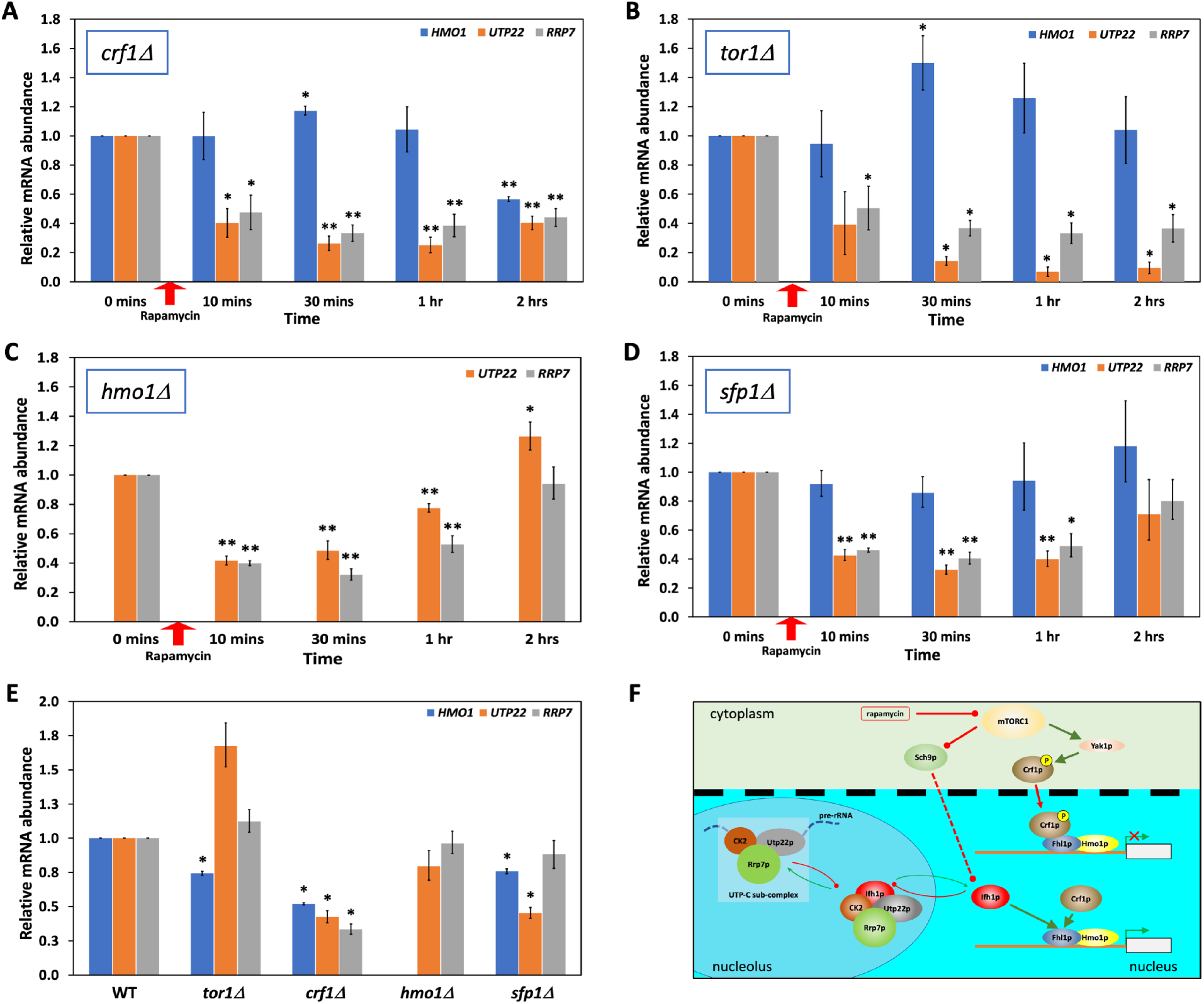
Gene expression in deletion strains. A. Relative abundance of *HMO1* (blue), *UTP22* (orange), and *RRP7* (grey) mRNA before and at the indicated times after addition of rapamycin to *crf1Δ* cells (red arrow). B. Relative mRNA abundance in *tor1Δ* cells on exposure to rapamycin. C. Relative mRNA abundance in *hmo1Δ* cells on exposure to rapamycin. D. Relative mRNA abundance in *sfp1Δ* cells on exposure to rapamycin. E. Relative mRNA abundance in unsupplemented cultures of the indicated strains relative to wild type cells; experiments with wild-type and deletion strains were done side-by-side. Error bars represent standard deviation from three biological replicates. For Panels A-D, transcript levels were calculated using 2^-ΔΔC^_T_ relative to the reference gene and reported relative to the corresponding unsupplemented cultures (0 min). Asterisks denote statistically significant differences between unsupplemented and rapamycin-supplemented cultures based on a Student’s t-test; *, p<0.05**; p<0.001. For Panel E, transcript levels were calculated using 2^-ΔΔC^_T_ relative to the reference gene and reported relative to wild-type cells. Asterisks denote statistically significant differences between wildtype and the indicated deletion strain based on a Student’s t-test; *, p<0.05. F. Proposed model for the role of Crf1p in gene regulation. When mTORC1 is active, target genes may be activated by binding either Ifh1p or Crf1p (bottom gene). On inhibition of mTORC1, activation of Yak1p leads to phosphorylation of cytoplasmically localized Crf1p and its translocation to the nucleus where it functions as a repressor of *HMO1* expression but not *UTP22* or *RRP7* despite binding the gene promoters (top gene). When mTORC1 is inhibited, Sch9p participates in removal of Ifh1p from gene promoters, causing repression of gene activity, and Ifh1p translocates to the nucleolus to generate the CURI complex in a process that requires Utp22p. As Crf1p is not as readily removed from gene promoters, repression as a result of mTORC1 inhibition may be mitigated.

### *TOR1* deletion resulted in the loss of *HMO1* but not *UTP22* and *RRP7* regulation

The decrease in *HMO1* mRNA abundance upon rapamycin treatment depends on Tor1p with no significant change in mRNA abundance after 1 h of rapamycin treatment of *tor1Δ* cells (Grove, 2018; Panday et al., 2017). Examination of mRNA levels at earlier time points revealed a significant ∼1.5-fold increase in *HMO1* mRNA 30 min after rapamycin addition, followed by a reduction to initial levels (Figure 4B). Notably, this transient increase in *HMO1* mRNA parallels that observed on deletion of *CRF1*. In contrast, levels of *UTP22* and *RRP7* mRNA were still reduced in *tor1Δ* cells. For *UTP22* and *RRP7*, mRNA levels were ∼10% and ∼35% of initial levels after 1-2 h of incubation of *tor1Δ* cells with rapamycin, which is similar to the changes observed in wild-type cells (Figures 1A and 4B). This suggests that Tor1p is critical for regulation of *HMO1* expression in response to mTORC1 inhibition, but not for regulation of *UTP22* and *RRP7* expression.

Using a strain expressing 3X-FLAG-tagged Tor1p, we previously reported the direct binding of Tor1p to the *HMO1* promoter and its dissociation upon prolonged (>1 h) rapamycin treatment. We performed ChIP on 3X-FLAG-tagged Tor1p and confirmed the expected Tor1p binding to the *HMO1* gene, centered near the IFHL site, and its dissociation after 2 h of rapamycin treatment. By contrast, there was no significant binding of Tor1p observed on either *UTP22* or *RRP7* genes before or after incubation with rapamycin (Figure S3). Combined with the differential requirement for Tor1p to achieve reduced gene expression on addition of rapamycin, we suggest that *HMO1* expression is controlled by direct promoter binding of Tor1p, whereas repression of *UTP22* and *RRP7* may be achieved indirectly by cytoplasmic mTORC1 containing either Tor1p or Tor2p.

Since Hmo1p and Fhl1p binding has been reported to be mutually dependent on some RP genes (Hall et al., 2006), we also examined gene expression in *hmo1Δ* cells. Absence of Hmo1p altered the time-dependent response to rapamycin, with an ∼60% reduction in both *UTP22* and *RRP7* mRNA levels after 10-30 min of rapamycin exposure (Figure 4C), as also observed in WT cells (Figure 1A). However, mRNA levels increased to original levels after prolonged rapamycin treatment, indicating that Hmo1p is required to maintain repression; such requirement for Hmo1p was not observed on RP genes (Figure S2D). Sfp1p has been implicated in mediating gene expression in response to mTORC1 activity, so we measured mRNA abundance in *sfp1Δ* cells. *HMO1* mRNA levels remained unchanged in *sfp1Δ* cells after addition of rapamycin, indicating that Sfp1p directly or indirectly participates in communicating mTORC1 inhibition at the *HMO1* gene (Figure 4D). By contrast, *UTP22* and *RRP7* mRNA abundance decreased on addition of rapamycin to *sfp1Δ* cells followed by an increase with prolonged rapamycin treatment, suggesting that Sfp1p is dispensable for the initial response to rapamycin but its absence interferes with maintenance of the repressed state. For the tested RP genes, absence of Sfp1p did not affect the response to rapamycin (Figure S2E).

### Crf1p functions as an activator

*HMO1* expression is modestly reduced in a *tor1Δ* strain during vigorous growth, indicating that Tor1p activates expression (Panday et al., 2017). In contrast, expression of *UTP22* and *RRP7* was unaffected by the absence of Tor1p (Figure 4E). Surprisingly, deletion of *CRF1* resulted in >2-fold lower levels of *HMO1, UTP22*, and *RRP7* mRNA. By comparison, expression of RP genes was not markedly altered in *tor1Δ* cells but expression of Class I and II RP genes, which have been reported to depend primarily on Fhl1p and its cofactors for regulation (Fermi et al., 2016; Knight et al., 2014; Zencir et al., 2020), was reduced on deletion of *CRF1* (Figure S2F). The reduction in mRNA levels in *crf1Δ* cells suggests that Crf1p functions as an activator. By contrast, deletion of *HMO1* had no significant effect on expression of either *UTP22* or *RRP7*, while deletion of *SFP1* led to reduced expression of *HMO1* and *UTP22*.

## Discussion

Yeast mTORC1 may be assembled with either of the paralogous kinases Tor1p or Tor2p, although Tor1p is preferred. Yeast mTORC1 is mainly activated by amino acid sufficiency; constitutively localized to the vacuolar membrane, activated mTORC1 in turn phosphorylates Sch9p, which promotes protein translation among other functions. The cellular localization of Tor1p and Tor2p is non-identical, consistent with a primary role for Tor2p as a component of mTORC2, and while both proteins have been reported to localize to the cytoplasm, only Tor1p has been detected in the nucleus (Li et al., 2006; Sturgill et al., 2008; Tsang et al., 2003). The observation that deletion of Tor1p has no effect on rapamycin-mediated reduction in *UTP22* and *RRP7* mRNA abundance implies that Tor2p can substitute for Tor1p; our data therefore indicate that rapamycin-mediated regulation of *UTP22* and *RRP7* by mTORC1 is indirect and may be mediated by cytoplasmically localized mTORC1. Sch9p, which colocalizes with mTORC1 to the vacuolar membrane, has been reported to be involved in the rapid release of Ifh1p from RP genes in response to mTORC1 inhibition, although Ifh1p is not known to be a direct target of Sch9p (Cai et al., 2013), and Sch9p may serve the same function at *UTP22* and *RRP7* (Figures 4F and S4). By contrast, since *HMO1* mRNA abundance was not reduced on addition of rapamycin to *tor1Δ* cells, we surmise that *HMO1* expression is controlled by nuclear (possibly promoter-bound) mTORC1 only. For *HMO1*, deletion of *TOR1* also resulted in reduced expression during balanced growth. This effect of *TOR1* deletion could also reflect that nuclear mTORC1 activity is required for phosphorylation of factors, which mediate active transcription. However, indirect effects cannot be ruled out as cytoplasmic mTORC1 activity is also reduced in *tor1Δ* cells as indicated by the observation that while calorie restriction or deletion of either *TOR1* or *SCH9* increases replicative life span, nutrient limitation in *tor1Δ* cells does not further increase life span (Kaeberlein et al., 2005).

While *TOR1* deletion has no effect on *UTP22* and *RRP7* mRNA abundance in response to rapamycin (Figure 4B), deletion of individual transcription factors affects gene regulation. Deletion of either *SFP1* or *HMO1* attenuates the response to rapamycin, particularly upon prolonged mTORC1 inhibition (Figure 4C-D), indicating that both Sfp1p and Hmo1p participate in communicating mTORC1 activity on the *UTP22* and *RRP7* genes. It should be noted, however, that deletion of *SFP1* results in mTORC1 activation, possibly on account of amino acid accumulation due to inhibition of translation (Lempiainen et al., 2009); such mTORC1 activation may in turn mitigate the effect of rapamycin addition. By contrast, deletion of *CRF1* is not required for repression of either *UTP22* or *RRP7* expression, analogous to the observation that Crf1p is not required for repression of RP gene expression on mTORC1 inhibition in the W303 yeast strain (Figures 4A and S2C) (Zhao et al., 2006).

For the *HMO1* promoter, absence of Hmo1p attenuates the response to rapamycin, comparable to the effect reported here on the *UTP22* and *RRP7* genes (Xiao et al., 2011). However, the rapamycin-mediated repression of *HMO1* expression appears to be completely dependent on both Sfp1p and Crf1p. *HMO1* expression is modestly reduced in *sfp1Δ* cells, consistent with Sfp1p functioning as an activator (Figure 4E). Sfp1p is directly phosphorylated by mTORC1, and inhibition of mTORC1 results in Sfp1p leaving the nucleus (Lempiainen et al., 2009). That Crf1p is also required for repression of *HMO1* expression on inhibition of mTORC1 is particularly notable as it indicates that the repressor function of Crf1p is not generally lost in this yeast strain, but that its effects are gene-specific and may depend on individual promoter architectures.

It was recently reported that overexpression of *CRF1* rescues the lethality of an *IFH1* deletion, suggesting that it can substitute for Ifh1p as an activator when in excess, and that it prevents rapamycin-mediated repression of Category I and II RP genes, which depend on Fhl1p and Ifh1p for regulation of gene expression (Zencir et al., 2020). Notably, our data indicate that Crf1p functions as an activator under normal growth conditions (Figures 4F and S4). By comparison to Ifh1p, Crf1p lacks N-and C-terminal regions implicated in removal of Ifh1p from RP gene promoters under conditions of mTORC1 inhibition (Albert et al., 2016). The absence of these extensions may therefore favor binding of Crf1p over Ifh1p. On genes where Crf1p acts as a repressor, whether the *HMO1* gene in W303 (Figure 4A) or RP genes in a strain such as TB50 (Martin et al., 2004), such preferred binding may lead to repression. However, our data suggest that preferred binding of Crf1p to specific genes may also occur during balanced growth, leading to activation of gene expression. While the binding and release of Ifh1p is a primary method by which gene expression may be controlled in response to mTORC1 inhibition, Crf1p may operate as a “fail-safe” activator to ensure adequate gene expression, even in the face of mTORC1 inhibition. Specifically, Hmo1p is intricately involved in regulation of both RNA Pol I and Pol II transcribed genes, and too efficient repression of *HMO1* may therefore be undesirable, an outcome that may be averted by using Crf1p as a primary regulatory transcription factor. We speculate that the ability of Crf1p to function as either an activator or a repressor is related to its phosphorylation state. Since Utp22p is required to prevent re-binding of Ifh1p to gene promoters under conditions of mTORC1 inhibition (Albert et al., 2016), too efficient repression of *UTP22* may likewise be detrimental.

Crf1p and Ifh1p derive from a single ancestral protein, which was proposed to function as an activator in rich media and as a repressor under condition of stress and nutrient limitation (Wapinski et al., 2010). Our data indicate that Crf1p encompasses the activator and repressor functions of the Ifh1p-Crf1p ancestor and that its functions depend on individual promoter architectures. This flexibility permits a sensitive regulation of genes encoding proteins that are central to the stress response.

## Acknowledgments

We thank J. Haber for providing plasmid pGAL-HO, A. Panday for helpful discussions and critical comments on the manuscript, and S. Herke and D. P. Singh for facilitating qPCR. This work was supported by the National Science Foundation (MCB-1714219 to A. Grove) and by a donation from P. Campbell in support of undergraduate research.

## Author Contributions

SK and AG, Conceptualization; SK, Methodology, Validation, Formal Analysis; SK, MM, PB, Investigation; SK, Writing – Original Draft; AG, Writing – Review and Editing; SK, Visualization; AG, Supervision, Funding Acquisition.

## STAR Methods

### RESOURCE AVAILABILITY

#### Lead Contact

Further information and requests for resources and reagents should be directed to and will be fulfilled by the Lead Contact, Anne Grove.

#### Materials Availability

Yeast strains are available upon request.

#### Data and Code Availability

This study did not generate or analyze datasets or code.

## EXPERIMENTAL MODEL AND SUBJECT DETAILS

Yeast cells were grown in YPD (yeast extract, peptone and 2% dextrose) or in the desired synthetic dropout media at 30°C in a shaking incubator at 200 rpm. Strain background is W303-1a.

## METHOD DETAILS

### Yeast strains and Plasmids

The W303-1a (*MAT****a*** *ADE2 his3-11 leu2-3, 112 lys2Δ trp1-1 ura3-1 can1-100*) isogenic yeast strain *DDY3* was used as a wild type strain. All other strains used were derived from *DDY3*. Strains expressing C-terminally FLAG-tagged Ifh1p, Crf1p, Hmo1p, Fhl1p, and Tor1p and the strain deleted for *TOR1* and *HMO1* were described previously (Panday *et al*. 2017, Xiao *et al*., 2011). The strain deleted for *CRF1* was created by amplification of the *LEU2* marker from pRS315 using primers having ∼80 nucleotide long flanking sequence homologous to the *CRF1* gene. Deletion of *SFP1* was accomplished similarly by amplifying the *TRP1* marker from pRS424 (Sikorski and Hieter, 1989). The resulting PCR products were transformed into *DDY3* to create *crf1Δ* and *sfp1Δ* strains. For primer sequences, see Table S1. Plasmid pGAL-HO expressing HO endonuclease under *GAL1* promoter was a gift from J. Haber (Brandeis University) (Herskowitz and Jensen, 1991).

### High-efficiency transformation

Cells were grown in YPD (yeast extract, peptone and 2% dextrose) at 30°C in a shaking incubator at 200 rpm and pelleted at OD_600_ ∼0.8. Pelleted cells were washed with 1X PBS (phosphate-buffered saline) and resuspended in 1X TEL (Tris, EDTA, and lithium acetate buffer) followed by overnight nutated incubation at room temperature. Cell pellets were collected and resuspended in 100 µl 1X TEL for each 10 ml of original culture and incubated for 30 mins at room temperature. Transformation of 100 µl of competent cells was performed with 10 µl of carrier DNA and 1 µg of plasmid DNA incubated at room temperature for 30 mins. Seven hundred µl of 40% PEG (polyethylene glycol) in 1X TEL was added to each transformation tube before incubation at room temperature for 1 hour without shaking. At the end of the 1hr incubation, 88 µl DMSO was added to each transformation tube followed by a heat shock at 42°C for 45 mins. Cells were pelleted at 8000 rpm for 30 seconds and washed with 300 µl water and resuspended in 400 µl water. Transformed cells were plated on desirable synthetic drop-out media.

### RNA isolation and in vivo gene expression

Yeast cells were grown to OD_600_ of 0.6 – 0.8 at 30°C. Cells were then treated with 200 nanograms of rapamycin (R0395; MilliporeSigma) dissolved in DMSO per milliliter of culture for analysis of mTORC1 inhibition. For inducing DNA damage, cells carrying plasmid coding for HO endonuclease under control of the *GAL* promoter were grown in synthetic dropout (SD) media with 2% w/v raffinose as a carbon source until OD_600_ 0.6-0.8. *MAT*-specific DSB was induced by adding 2% galactose to the media to express *HO*. DNA damage induction was terminated by adding 2% glucose. Replication stress caused by dNTP depletion was created by treating cells grown in YP (yeast peptone) with 0.2 M final concentration of hydroxyurea (H8627; MilliporeSigma).

A total of 5 mL culture was harvested prior to treatments (for untreated control) and at the indicated time points. Cells were pelleted and washed with ice-cold diethyl pyrocarbonate (DEPC) treated water and pelleted again. Pellets were frozen at −80°C after discarding the supernatant. Total RNA was extracted using illustra RNAspin Mini isolation kit (Cytiva) or Monarch Total RNA Miniprep Kit (New England Biolabs) subsequent to spheroblast construction using 10 U of Zymolyase (ZYMO Research) per sample for 30 mins at 37°C. Total RNA extracted was rendered free of any genomic DNA contamination using Turbo DNase (Invitrogen) and absence of DNA was verified by PCR. RNA quantification was done using NanoDrop (Thermo Scientific) and all sample concentrations were normalized to 100 ng/µl by adding the required volume of nuclease free water. The mRNA abundance levels were determined using 200 ng total RNA for each sample in technical replicates using Luna Universal One-Step RT-qPCR kit (New England BioLabs) and SYBR Green as detection agent and gene-specific primers (Table S2). All quantitative PCR reactions were performed using QuantStudio-6/ViiA7/7900HT Fast Real-Time PCR System (Applied Biosystems) using 96-well or 384-well plates. Data analysis was performed after normalization of expression levels using *IPP1* as a reference gene for rapamycin treatment or induction of DNA damage, while expression of *ACT1* was used as a reference control for HU treatment. All mRNA abundance levels were calculated using the ΔΔC_t_ method and experiments were repeated at least three times (biological replicates). Reported average, standard deviation and statistical significance were calculated using Student’s t-test.

### ChIP and quantitative PCR

Yeast cells expressing FLAG-tagged proteins of interest were grown in YPD until the OD_600_ reached ∼1.0. A 50 ml aliquot was collected before treatment to serve as untreated control and the remaining culture was treated with a final concentration of 200 ng/mL rapamycin. Volumes were adjusted to an equivalent of OD_600_ = 1.0 for aliquots collected for subsequent time points post rapamycin treatment. Aliquots of the culture were immediately fixed using 1% formaldehyde final concentration at room temperature by gentle shaking continuously for 20 minutes. Formaldehyde crosslinking was stopped by addition of an equivalent volume of 2.5 M glycine as the volume of formaldehyde. Cells were pelleted and washed twice using 1X PBS and frozen at −80°C after discarding the supernatant.

Cells were mechanically lysed by vortexing with 0.5 mm glass beads for 40 mins at 4°C in 400 µl lysis buffer (50 mM HEPES, pH 7.5, 140 mM NaCl, 1% Triton X-100, and 1% sodium deoxycholate) with 10 µl/ mL of protease inhibitors prepared from Roche-cOmplete, EDTA free Protease inhibitor cocktail tablets (1 tablet/mL nuclease free water). Ten µl phenylmethylsulfonyl fluoride (PMSF; 100 µM) was added to each sample before and after chromatin shearing to prevent protease activity. Chromatin shearing was achieved by sonicating the cell lysates 6 times for 10 seconds each round at 25% amplitude resulting in fragment size peaking at ∼300 base pairs as determined by agarose gel electrophoresis. Cell lysates were chilled on ice for at least one minute between each round of sonication to prevent heat induced protein denaturation. Each sample was divided in three 100 µl aliquots for input control, immunoprecipitation, and no antibody control. Cell lysates were precleared for immunoprecipitation and no antibody using protein G-Sepharose beads (Cytiva) to reduce background signals from nonspecific binding. Immunoprecipitation was performed with 5 µg monoclonal anti-FLAG antibody (MilliporeSigma; F1804). Eluted ChIP-DNA samples were analyzed by 30 cycle PCR amplifications followed by electrophoresis on 1.5 % agarose gels and visualization with ethidium bromide and by quantitative real time PCR. The qPCR was performed with Luna Universal qPCR Master Mix (New England BioLabs) using SYBR Green detection. All qPCR runs were performed using QuantStudio-6/ ViiA7/ 7900HT Fast Real-Time PCR System (Applied Biosystems) on 96-well or 384-well plates. Data analysis was performed following normalization with untreated control for time points indicated after treatment. The percentage of IP compared to input control of each time point was calculated after subtracting background signals. Three primer pairs each for *HMO1, UTP22*, and *RRP7* were used to detect protein binding at different regions (Table S3). Average values, standard deviations and significance values calculated using Student’s t-test were obtained and reported from three independent biological replicates.

## QUANTIFICATION AND STATISTICAL ANALYSIS

Statistical analyses were performed using Student’s t-test in Excel. In all cases, n represents at least 3 biological replicates and data are represented as mean ± SD. Statistical significance is defined as p < 0.05. Details may be found in figure legends.

