## Supplemental information for "Intricate regulation of ribosome biogenesis genes in response to mTORC1 signaling"

Contents:

Figure S1. Transcription factor binding to the *HMO1* gene. Related to Figures 2 and 3.
Figure S2. Repression of RP genes in response to rapamycin. Related to Figure 4.
Figure S3. Tor1p binds directly to *HMO1* but not to *UTP22* or *RRP7.* Related to Figure 4.
Figure S4. Proposed model for the role of Crfp in gene regulation. Related to Figure 4.
Table S1. Primers used for strain construction. Related to Figure 4.
Table S2. Primers used for qRT-PCR. Related to Figures 1 and 4.
Table S3. Primers used for ChIP. Related to Figures 2 and 3.

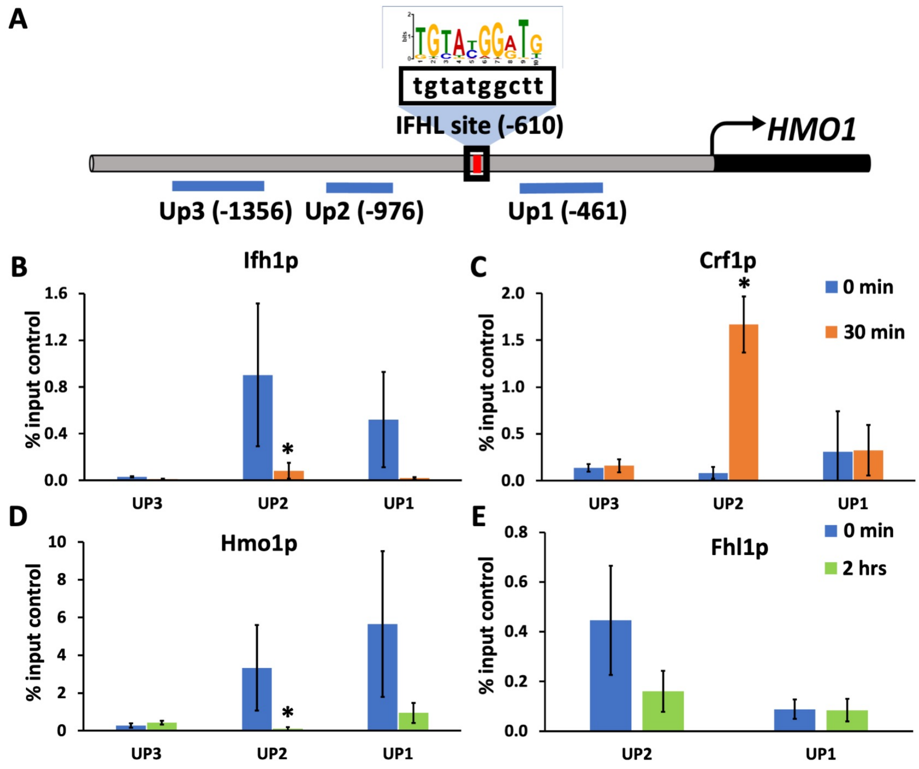


**Figure S1. Transcription factor binding to the *HMO1* gene. Related to Figures 2 and 3.** A. *HMO1* upstream region with IFHL site shown in red and expanded to show sequence; consensus IFHL site shown above. Positions of amplicons are identified with numbers representing the upstream edge. B-E. Binding of Ifh1p, Crf1p, Hmo1p, and Fhl1p at the indicated positions as determined by ChIP using antibodies to FLAG-tagged proteins. Blue bars represent binding before addition of rapamycin, while orange and green bars represent binding detected 30 min and 2 h following addition of rapamycin, respectively. Data are normalized to the corresponding input control and are presented as the average of three biological replicates; error bars represent SD. Asterisks denote statistically significant differences before (0 min) and after addition of rapamycin based on a Student's t-test; *, $p<0.05$.

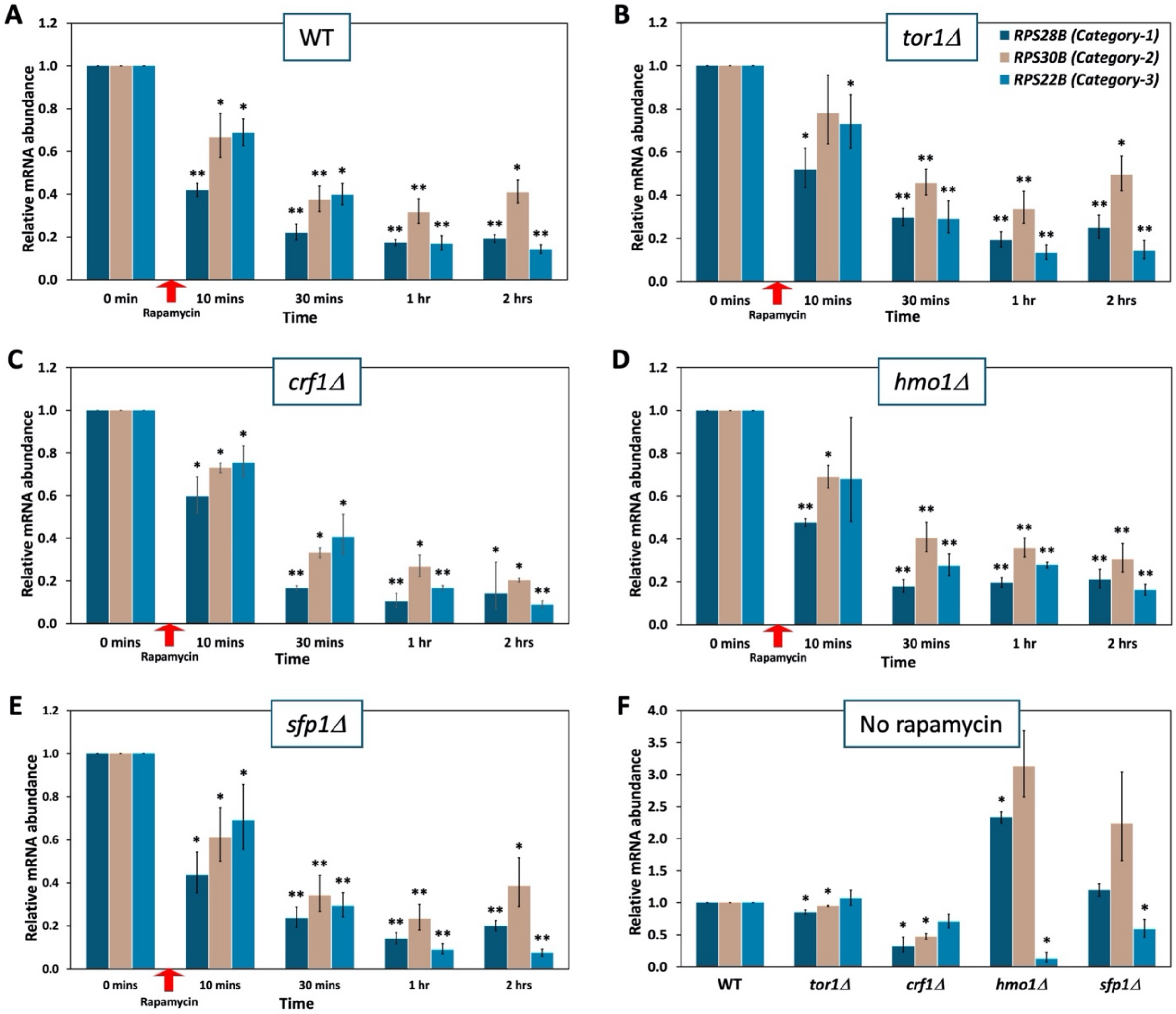


**Figure S2. Repression of RP genes in response to rapamycin. Related to Figure 4**. Relative abundance of *RPS28B* (deep teal), *RPS30B* (wheat), and *RPS22B* (teal) mRNA (legend as inset to Panel B). The identified genes are examples of each of three categories of RP genes, with genes belonging to categories I and II binding transcription factors Rap1p, Sfp1p, Fhl1p, and Ifh1p and Category I genes in addition binding Hmo1p. Thirteen genes belong to Category III, whose regulation mainly depends on the general transcription factor Abf1 and Sfp1p. Relative mRNA abundance is reported at the indicated times after administration of rapamycin to wild-type (A), *torΔ* (B), *crf1Δ* (C), *hmo1Δ* (D), and *sfp1Δ* (E) cells. Transcript levels were calculated using $2^{-\Delta\Delta C_T}$ relative to the reference gene and are reported relative to the corresponding unsupplemented cultures (0 min). F. Relative mRNA abundance in unsupplemented cultures of the indicated strains relative to wild type cells. Error bars represent standard deviation from four biological replicates. Asterisks denote statistically significant differences between unsupplemented and rapamycin-supplemented cultures (Panels A-E) or between wildtype and the indicated deletion strain (Panel F) based on a Student's t-test; *, p<0.05; **, p<0.001.

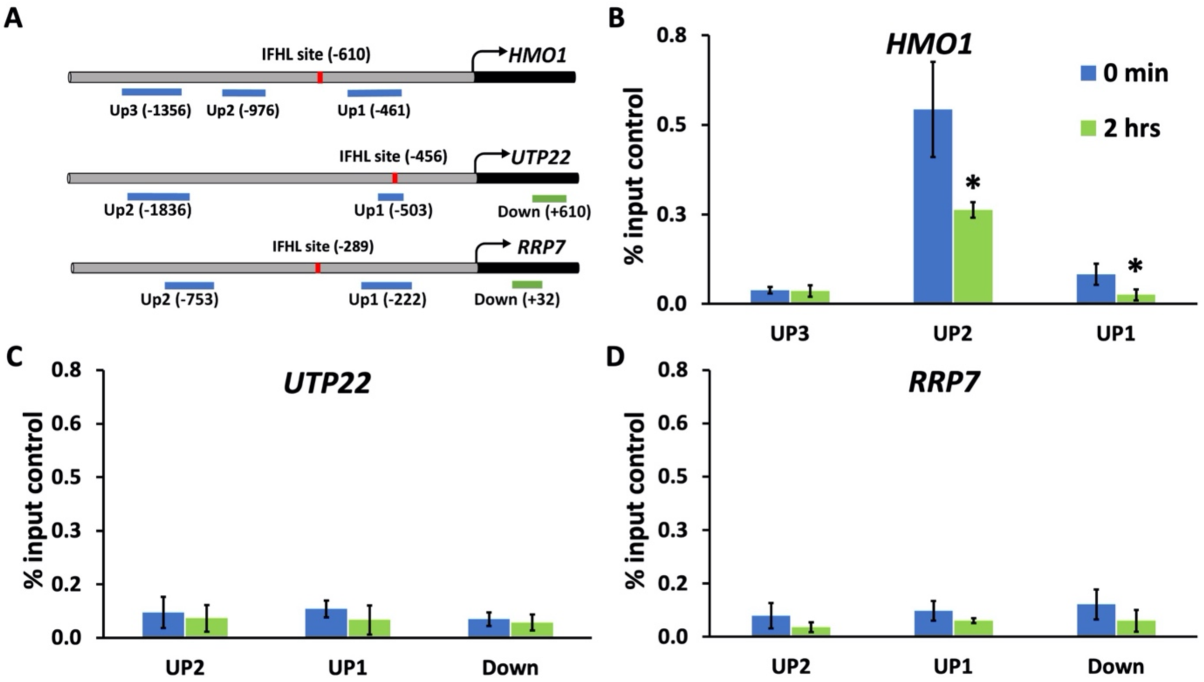


**Figure S3. Tor1p binds directly to *HMO1* but not to *UTP22* or *RRP7*. Related to Figure 4.** A. Position of amplicons and IFHL sites (confirmed for *HMO1*, predicted for *UTP22* and *RRP7*). B-D. Binding of Tor1p at the indicated positions as determined by ChIP using antibodies to FLAG-tagged Tor1p. Blue bars represent binding before addition of rapamycin, while green bars represent binding detected 2 h following addition of rapamycin. Data are normalized to the corresponding input control and are presented as the average of three biological replicates; error bars represent SD. Asterisks denote statistically significant differences before (0 min) and after addition of rapamycin (2 hrs) based on a Student's t-test; *, $p<0.05$.

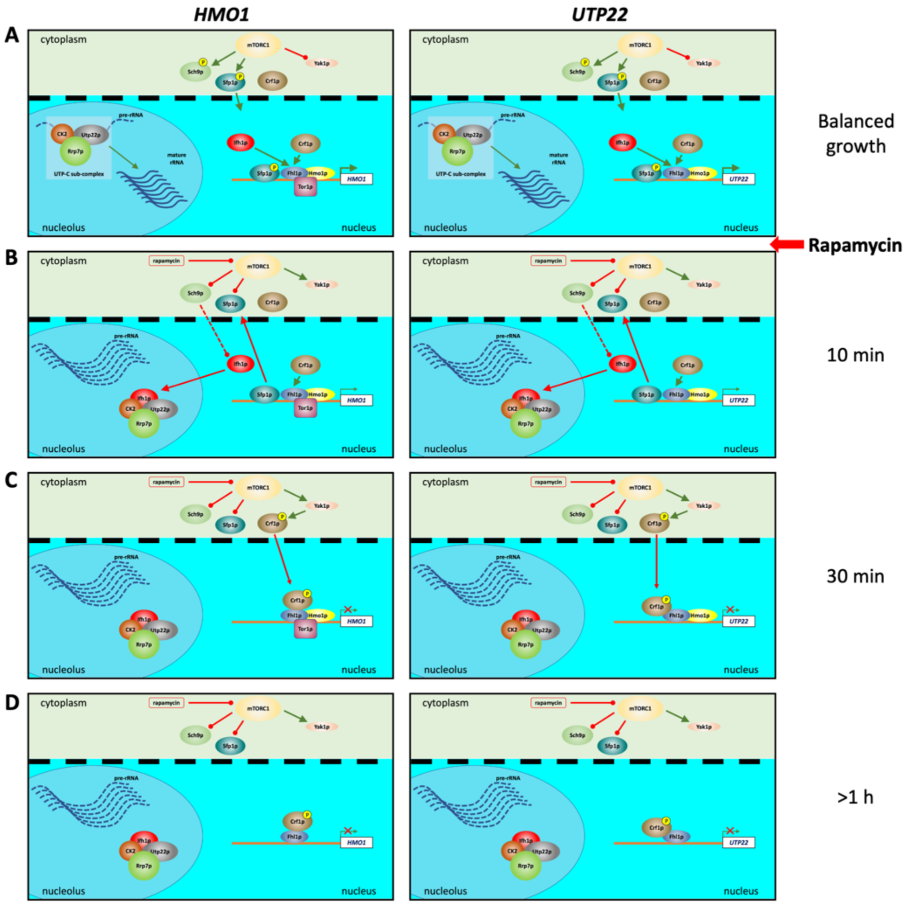


**Figure S4. Proposed model for the role of Crfp in gene regulation. Related to Figure 4**. A. When mTORC1 is active, direct targets Sch9p and Sfp1p are phosphorylated, which promotes translation, transcription of genes encoding components and assembly factors of ribosomes, and pre-rRNA processing. Either Ifh1p or Crf1p function to activate transcription. B. Inhibition of mTORC1 by addition of rapamycin results in rapid dissociation of Ifh1p in a process that requires Utp22p. This generates the CURI complex, resulting in suppression of both pre-rRNA processing and transcription of Ifh1p target genes. Sfp1p also rapidly leaves the nucleus. C. Activation of Yak1p under conditions of mTORC1 inhibition ultimately leads to phosphorylation of cytoplasmically localized Crf1p, which translocates to the nucleus. While Crf1p binds both *HMO1*, *UTP22*, and *RRP7* genes, it is required as a repressor only for *HMO1* (left). D. Prolonged exposure to rapamycin results in dissociation of Tor1p from the *HMO1* gene as well as dissociation of Hmo1p.

**Table S1. Primers used for strain construction. Related to Figure 4.**

| Primer name | Sequence (5' to 3') |
|---|---|
| Crf1Forward | GCGGTCACTAGAAAGTTATAAGAAGGTTACCATTTAACTGAACAGCAGCGGTATAGTACGGAATAAAAAAGTCTTTTAAAGCAGATTGTACTGAGAGTGC |
| Crf1Reverse | AGTAAAGTGCTTCTATTTTTTCGCAAACGATCATATTTTCCTTTAAAAGACGATAATGATTCTTCTGGAATATCAAATAACTCCTTACGCATCTGTGCGG |
| Sfp1Forward | TTCAAATTTATTGTTATTAAAAGACATATTCATATTCAGTATTTTTATTTTTCTTCGAAACAGGAAAAGCTTATCTGAGTGCAGATTGTACTGAGAGTGC |
| Sfp1Reverse | GCTCGATGCATGAGTCGGTAACGGTAGAATTAGTATCTAAATAGCATACTAAATCAATTCAGTAAAGAAACAATTATATCCTCCTTACGCATCTGTGCGG |

**Table S2. Primers used for qRT-PCR. Related to Figures 1 and 4.**

| Primer name | Sequence (5' to 3') |
|---|---|
| IPP1F | CCCAATCATCCAAGACACCAAGAAGG |
| IPP1R | AGCAATAGTTTCACCAATTTCCAACACATC |
| HMO1F | GCTCCAGTCAAGGCTGTAAG |
| HMO1R | CACGAAGTTCTTGACGAACG |
| UTP22F | TGCTTACACAAGAGGAGCGT |
| UTP22R | TCACCCGTTGGTTTGGAACA |
| RRP7F | ACGGGTTTATAGTGGTGCCG |
| RRP7R | TATTTCCCACAGAGCTGCCC |
| RPS30B_F | CTCTAGCTCGTGCTGGTAAAGT |
| RPS30B_R | GCACGACCCTTTGGTTTCTT |
| RPS28B_F | GGAAGACACTTCCCGTACCA |
| RPS28B_R | TTAACGCAAACGACGAGCTTC |
| RPS22B_F | AGACCGGTAAACGTCAGGTTC |
| RPS22B_R | GCAATAAGTTGGCGGTCCAT |

**Table S3. Primers used for ChIP. Related to Figures 2 and 3.**

| Primer name | Sequence (5' to 3') |
|---|---|
| HMO1 Up1F | CTTAGGCACACGTATGCACATATCCA |
| HMO1 Up1R | GTGGTACTGGAGAGAATCTGAGTGAA |
| HMO1 Up2F | TCCTGAAACCCAAAGATCCTGAAGTC |
| HMO1 Up2R | AGATGAGCGGTTAAGTGATTCCACTG |
| HMO1 Up3F | ATCATCATCGTCGTCATCATCGTCTA |
| HMO1 Up3R | GACTTCAGGATCTTTGGGTTTCAGGA |
| UTP22 DownF | TGCTTACACAAGAGGAGCGT |
| UTP22 DownR | TCACCCGTTGGTTTGGAACA |
| UTP22 Up1F | GGAAGTTCTGTCCATCCGGT |
| UTP22 Up1R | CACCGGTCCTGCCTAAATCT |
| UTP22 Up2F | ATTGGCGAAGAGATGGAGGC |
| UTP22 Up2R | ATTGGCTAGTGTTCACGGCA |
| RRP7 Down1 | ACGGGTTTATAGTGGTGCCG |
| RRP7 DownR | TATTTCCCACAGAGCTGCCC |
| RRP7 Up1F | AGAGGTAGTAGCAAGAGTGGGT |
| RRP7 Up1R | TTCGTTCGCCCTAAATGCCT |
| RRP7 Up2F | TGACGGTTACGAAGAAGCCC |
| RRP7 Up2R | TCGTTAGTGTTCGGTTTCCAAG |
